# Capturing variation in metagenomic assembly graphs with MetaCortex

**DOI:** 10.1101/2021.07.23.453484

**Authors:** Samuel Martin, Martin Ayling, Livia Patrono, Mario Caccamo, Pablo Murcia, Richard M. Leggett

## Abstract

**Motivation:** The assembly of contiguous sequence from metagenomic samples presents a particular challenge, due to the presence of multiple species, often closely related, at varying levels of abundance. Capturing diversity within species, for example viral haplotypes, or bacterial strain-level diversity, is even more challenging.

**Results:** We present MetaCortex, a metagenome assembler that captures intra-species diversity by searching for signatures of local variation along assembled sequences in the underlying assembly graph and outputting these sequences in sequence graph format. We show that MetaCortex produces accurate assemblies with higher genome coverage and contiguity than other popular metagenomic assemblers on mock viral communities with high levels of strain level diversity, and on simulated communities containing simulated strains.

**Availability and Implementation:** Source code is freely available to download from https://github.com/SR-Martin/metacortex, is implemented in C and supported on MacOS and Linux.

**Supplementary information:** Supplementary materials are available at the journal’s website. All assemblies, simulated reads, and simulated genomes used in this paper have been deposited online on Zenodo and can be found at DOI 10.5281/zenodo.6616437.

## INTRODUCTION

The well documented increase in yield and reduction in cost of DNA sequencing technologies has led to a rapid increase in the use of shotgun approaches for studying metagenomic samples [1]. A first analysis step is often taxonomic classification of reads by comparison with reference databases. However, deeper analysis is enabled by assembling sequence data to form longer contiguous sequence (contigs). Such assembly may facilitate improved classification, clustering of sequences (particularly where reference genomes are unavailable), or analysis at the scale of genes.

There are two fundamental approaches for the assembly of sequencing data: Overlap-layout-consensus (OLC) assembly, and de Bruijn graph assembly. In OLC assembly, each read is compared to every other read, and reads that overlap well are merged together to form contigs. The de Bruijn graph technique utilises directed graphs to represent the *k*-mers (short sequences of length *k*) present in a set of reads, and this representation of the read set turns the assembly problem into a graph traversal problem. Traversing a graph has a lower order of time complexity than OLC, and so the computational time to perform an assembly can be significantly reduced. Furthermore, the amount of memory required to build the graph is proportional to the total *k*-mers present in the sample, rather than the total number of reads. This is particularly important as new generations of sequencing technologies are producing ever greater amounts of data.

Metagenomic assembly presents several challenges beyond those of de novo genomic assembly [2]. The main additional difficulties are unknown diversity and unknown species abundance. Diverse species may not share *k*-mers, particularly if *k* is large, and in this case it can be inferred that subgraphs representing distinct species are mostly disconnected and existing assembly techniques from de novo genome assembly may be used on each disconnected subgraph. However, it is often not the case that a metagenomic sample consists solely of diverse species, and separating closely related species, or many strains of a single species, is a difficult task, particularly at low abundance.

The problem of capturing diversity below the species level in an assembly presents an even bigger challenge. The genomes of different strains from a single species can differ by single nucleotide polymorphisms (SNPs), large-scale structural variation, and anything in between. In many ways, the challenges faced here are similar to those faced when separating haplotypes in de novo assembly, where much of the same sequence is shared between haplotypes. In this case, one can use the ploidy of the organism and the *k*-mer coverages to guide the assembly. However, in the metagenomic case we do not necessarily know what the expected coverage of each strain is, and in low abundance cases it will be difficult to distinguish a SNP from a sequencing error. The challenge is at its most stark when studying viral metagenomes. Due to their short replication times, large population sizes and lack of proofreading mechanisms (Coronaviruses are a notable exception [4]), viruses can evolve extremely rapidly. As such, viruses are often referred to as a quasispecies consisting of a set of related strains; and the genome sequence of a strain is sometimes referred to as a haplotype [5].

Due to the throughput of current generation sequencing technologies, it is possible to have a metagenomic data set consisting of several terabytes of reads. Many assemblers are incapable of assembling such large datasets within realistic time and memory constraints. A common strategy to make the problem tractable is to randomly subsample the read set to obtain a much smaller one which can be assembled. This process will usually not affect the assemblies of high abundance species in the sample, but there is evidence that the assemblies of the low abundance species will be incomplete and of poorer quality [3].

Many current short-read metagenomic assemblers utilise the de Bruijn graph paradigm. Popular examples include Ray Meta [6], MEGAHIT [7], MetaVelvet [8], and metaSPAdes [9]. A key part of the implementation of all these tools is to collapse an assembly graph into linear sequences (usually in the form of a FASTA file). This facilitates easy downstream analysis, but the act of converting the assembly graph into linear sequence has the effect of removing the understanding of sequence diversity that is implicit in the graph. Several assemblers have been created specifically for the assembly of viral quasispecies, such as SAVAGE [10] and VICUNA [11]. These tools are able to assemble the haplotypes present in an isolated sample but may be less effective at assembling quasi-species from metagenomic samples. Within the (single-) genome assembly world, there is a growing awareness of the importance of capturing the genome graph in the output from assembly tools. This has resulted in the development of the FASTG format [12] and, more recently and with wider adoption, the graphical fragment assembly (GFA) format for assembly graph files [13]. These have been implemented in a number of tools including recent versions of ABySS [14], SPAdes [15] and SDG [16]. However, as yet, specialist metagenomic assembly tools lack the ability to capture variation through GFA output.

Here we introduce MetaCortex, a de Bruijn graph metagenomic assembler that is built upon data structures developed for the Cortex assembler [17]. As well as performing metagenomic assembly with standard FASTA output, MetaCortex generates assembly graph files that preserve intra-species variation (e.g. viral haplotypes), and implements a new graph traversal algorithm to output variant contig sequences. Whilst MetaCortex can be used to assemble any metagenomic dataset, we have developed features to specifically target metagenomic datasets with high levels of strain diversity (e.g. viral communities), and to represent this diversity in the resulting assembly. MetaCortex captures variation by looking for signatures of polymorphisms in the de Bruijn graph constructed from the reads and represents this in sequence graph format (both FASTG and GFA v2), and the usual FASTA format. The sequence graph provides information on local variation, such as SNPs and indels, along each contig identified by MetaCortex. By using the efficient data structures from Cortex, MetaCortex is capable of utilizing all *k*-mers from large metagenomic datasets, and able to perform assemblies from these datasets on a single CPU.

We show that MetaCortex is able to produce highly contiguous assemblies capturing almost all genome level diversity, and with a low level of misassemblies. By outputting sequence graph files, we were able to capture strain level diversity that is not present in the contigs and use this to manually assemble contigs that were specific to individual strains in a sample.

## METHODS

To test MetaCortex, we assembled real Illumina read sets from two mock communities of twelve viruses, at varying levels of abundance; a real human gut sample which was subsequently *in silico* mixed with real reads from a lab mix of five strains of HIV in equal abundance; and two simulated read sets from communities with high levels of strain variation. In each case we assembled the dataset with *k*-mer values equal to 31, 63, 95, and 127.

To compare MetaCortex’s performance with that of existing de-novo metagenome assemblers, we also assembled the same datasets using MEGAHIT (v1.1.1), Ray Meta (v2.3.1), MetaVelvet (v1.2.02), metaSPAdes (v3.14), and IDBA-UD (v1.1.2) [30]. Since MEGAHIT uses several *k*-mer values when constructing an assembly, we varied the --min-count parameter (with values default, 5, 10, and 20) for each dataset. For Ray Meta we used *k*-mer values equal to 31, 63, 95, and 127. For the real sequence data, we used trim-galore v0.5.0 [18] (a wrapper script around Cutadapt [19]), to trim low quality bases and adapter sequence from the reads, with the flags --paired and -- retain_unpaired where appropriate. Since this can break the pairing of paired end reads and result in single-ended reads, we assembled these sets using both “paired + unpaired single” and “single only” modes, for the assemblers where this was possible. The assemblies by IDBA-UD either failed, or were still running after 62 days, so the results for this assembler are not provided.

Statistics on the assemblies were obtained using MetaQUAST [20], which by default considers only those contigs greater than 500bp in length. A position on a contig is considered a misassembly by MetaQUAST if either the left flanking sequence aligns over 1kbp away from the right flanking sequence on a reference genome; flanking sequences overlap on more than 1kbp; or flanking sequences align to different strands, chromosomes, or genomes. We set the flag “ambiguity-usage” equal to one, so that only the best alignment from each contig is used when calculating certain statistics. Here, for each dataset we present a single assembly from each assembler, that we judged to be the best (based on genome coverage and error rates). Full results for all assemblies are available in the supplementary materials, along with the commands used.

### Assembly of sequenced mock viral communities

For our first benchmark, we assessed how well MetaCortex performs metagenomic assembly on simple mock communities compared to other current assemblers. Two mock communities of twelve viruses, each containing two ssDNA viruses and ten dsDNA viruses, were assembled from real Illumina read data made available for benchmarking purposes [21].

For both mocks, IDBA-UD was still running after 60 days, with 8 CPUs assigned. The assemblies by MetaVelvet were either killed by the Linux Out of Memory (OOM) killer after running out of memory, with 3TB of RAM allocated, or they recovered an insignificant total genome fraction, and so are not reported here. For MockB, metaSPAdes failed after running out of memory, with 3TB of RAM allocated. This is beyond the memory limits for many researchers, and the usual strategy at this point is to assemble a much smaller subsample of reads instead. However, recent studies suggest that subsampling can drastically reduce the length and proportion of conserved genes in the subsequent assembly, when compared to the assembly of the full dataset [3].

The first community, Mock A, was composed of the dsDNA viruses each (theoretically) at 9.82% abundance, and the ssDNA viruses at 0.92%. The read set consisted of 2×250bp paired end reads, sequenced on the Illumina MiSeq platform, with a total read count of 96m, including reads from host DNA. Using MetaCortex (SW, *k*=95), we were able to assemble 99.89% of the viral genomes with no misassemblies. Mismatch and indel rates were very low, at 4.43 per 100kbp and 3.44 per 100kbp respectively. Individual genome coverages ranged from 100% to 99.56%. Eight virus genomes were each assembled in a single contig, while the assembly of all other genomes ranged from 8 to 40 contigs. Figure 3.B shows that the assembly by MetaCortex is the most contiguous.

Table 1 shows that only the assembly by metaSPAdes recovers a higher genome fraction than the assembly by MetaCortex, with an extra 0.012%. However, metaSPAdes recovered only four species at 100% (compared with six for MetaCortex), and seven in a single contig (compared with eight for MetaCortex). The assembly by MetaCortex was the most contiguous (Figure 3.B) and has error rates almost identical to metaSPAdes (which had the lowest), with only a slightly higher indel rate. The assembly by MEGAHIT recovered a similar genome fraction across most species in the mock (Figure 3.A), but the assembly was less contiguous (Figure 3.B) and had the highest mismatch rate and the most misassemblies. Notably, the assembly by Ray Meta failed to assemble any of the genome of the ssDNA phage alpha3.

**Table 1:** Summary statistics for assemblies of Mock A. The contigs field describes the number of contigs in the assembly of length greater than or equal to 500bp. Mismatch and indel rates are the number of occurrences per 100kbp.

|  | Genome Fraction | Genomes Covered >50% | Contigs | Misassemblies | Mismatch rate | Indel rate |
| --- | --- | --- | --- | --- | --- | --- |
| MetaCortex | 99.886% | 12/12 | 339 | 0 | 4.43 | 3.44 |
| metaSPAdes | 99.898% | 12/12 | 2,259 | 0 | 4.43 | 3.12 |
| MEGAHIT | 98.233% | 11/12 | 1,902 | 3 | 75.7 | 3.33 |
| Ray Meta | 97.652 | 11/12 | 1,737 | 0 | 9.06 | 3.02 |

The total aligned length of the assembly by MetaCortex was 1.09Mbp, out of a total assembly length of 5.7Mbp. We used BLAST to query the longest unaligned contigs (greater than 100kbp) against the nt database. This revealed that each had alignments to one of Pseudoalteromonas or *Cellulophaga baltica*. These were host species that the mock community was grown on, so this represents legitimate assembly of DNA present in the sample. Including these species in the reference list for MetaQUAST, we had a total aligned length of 4.96Mbp, of which 3.87Mbp aligned to *C. baltica* over 143 contigs (covering 82.03% of the *C. baltica* genome).

The second community, Mock B, consisted of the dsDNA viruses at 3.51% abundance, and the two ssDNA viruses each at 32.47% abundance. The read set consisted of 98m 2×250bp paired end reads. Using MetaCortex (SW, *k*=63), we assembled 99.71% of the community, with a single misassembly (Table 2). The percent of individual genomes recovered ranged from 100% genome coverage, while the others ranged from 100% (for 5 genomes) to 98.744%. Seven virus genomes were assembled in a single contig, and the assembly of all other genomes varied from between six and 25 contigs. As was the case in Mock A, approximately 3.80Mbp of unaligned contigs were found to align to *C. baltica*, covering 81.804% of its genome.

**Table 2:** Summary statistics for assemblies of Mock B.

|  | Genome Fraction | Genomes Covered >50% | Contigs | Misassemblies | Mismatch rate | Indel rate |
| --- | --- | --- | --- | --- | --- | --- |
| MetaCortex | 99.711% | 12/12 | 735 | 1 | 7.23 | 3.45 |
| MEGAHIT | 94.15% | 11/12 | 1,762 | 2 | 15.66 | 3.65 |
| Ray Meta | 95.582% | 12/12 | 2,240 | 0 | 4.63 | 2.91 |

MetaCortex recovered the largest genome fraction, with a similar fraction recovered to Mock A. The assembly by MetaCortex was also the most contiguous (Figure 3.b), although this time contained a single misassembly (one more than Ray Meta) and had error rates between those in the assemblies by Ray Meta and MEGAHIT (Table 2).

### Assembly of sequenced HIV lab mix

In order to evaluate MetaCortex’s ability to capture variants in a real metagenomic sample, we created a read set containing five well-studied strains of HIV-1 (89.6, HXB2, JR-CSF, NL4-3, YU2) in equal abundance, that was bioinformatically mixed with reads from a human preterm baby gut sample. Reads and reference sequences for each HIV strain were made available in [22] (SRA run SRR961514). Each strain was between 93% and 97% identical to the other strains, and the read set consisted of 308Mb of Illumina 2×250bp reads. Reads from the human gut sample were taken from [27] (ENA run accession ERR2099157), consisting of 10.5Gb of Illumina 2×250bp reads. After mixing, reads from the HIV mix consisted of about 2.8%, representing about 0.56% per strain.

To increase the sensitivity of MetaCortex’s ability to capture strain level variation, we set the SW delta parameter to 0.4, and the min coverage parameter to 25. With these values, MetaCortex (SW, *k*=127) assembled 82.563% of the five HIV genomes across 31 contigs with no misassemblies.

We also assembled this dataset with MEGAHIT, Ray Meta, metaSPAdes, and MetaVelvet. Each of these tools produced assemblies with genome fractions below that produced by MetaCortex (Figure 4) and in some cases with higher misassembly rates (Table 3). Only Ray Meta produced assemblies with a significantly lower mismatch rate, but this assembly contained more misassemblies and a smaller genome fraction. Another assembly by Ray Meta (see supplementary materials) contained a smaller genome fraction (70.089%) with no misassemblies but higher mismatch and indel rates (76.67 and 11.8 respectively).

**Table 3:** Summary statistics for assemblies of 5-strain HIV mix.

| | Genome Fraction | Contigs ($\geq 500$bp) | HIV-Aligned Contigs | Misassemblies | Mismatch Rate | Indel Rate |
| --- | --- | --- | --- | --- | --- | --- |
| MetaCortex | 82.563% | 4,635 | 31 | 0 | 508.2 | 10.01 |
| MEGAHIT | 36.153% | 5,075 | 43 | 1 | 2744.27 | 68.61 |
| metaSPAdes | 36.153% | 3,409 | 10 | 0 | 1424.3 | 0 |
| MetaVelvet | 2.011% | 3,661 | 3 | 0 | 1747.17 | 0 |
| Ray Meta | 72.597% | 1,243 | 43 | 1 | 74.03 | 8.54 |

### GFA output captures strain level variation and facilitates visualisation

We used MetaCortex to create a sequence graph of the assembly, as an alternative method to capture strain-level diversity. Using the MC algorithm, we assembled the mixed reads and used BLAST [29] to map each contig against a database consisting of the reference genomes for each HIV strain. We found one contig of length 9,166bp (about the length of the HIV genome) that mapped well to the reference genomes. We extracted this contig, and the corresponding sequence graph elements into new files. Figure 4.C shows the resulting sequence graph, as visualised by GfaViz [23].

To demonstrate that this sequence graph contains more information than the corresponding contig, we constructed one contig for each of the five strains present in the sample. First, we created a mapping of each sequence in the graph to a database of the reference genomes using BLAST. Then, we constructed contigs in the following way. For each strain, a walk is performed through the sequence graph, where at each branch, the branch whose sequence has the highest score for the strain is chosen. Scores are calculated as the mapping identity multiplied by the proportion of the alignment covering the query (in ambiguous cases, the branch corresponding to the original contig is chosen). Each walk corresponds to a contiguous sequence, which forms the strain-level contig. A python script to parse the GFA file and BLAST mapping and construct the individual strain assemblies is available in the MetaCortex repository, under scripts/strain_assembly.py.

Next, we used dnadiff (part of MUMmer [24]) to compare each strain-specific contig and the original contig to the strain’s reference sequence. We found that the strain-specific contigs had alignments with a higher average identity to the corresponding reference strain and contained far fewer SNPs and indels (Table 4).

**Table 4:** Summary statistics comparing each strain-specific assembly to the base assembly.

|  | NL4-3 |  | JR-CSF |  | HXB2 |  | 89.6 |  | YU2 |  |
| --- | --- | --- | --- | --- | --- | --- | --- | --- | --- | --- |
|  | Original | Strain specific | Original | Strain specific | Original | Strain specific | Original | Strain specific | Original | Strain specific |
| Total Length | 9,166 | 9,125 | 9,166 | 9,166 | 9,166 | 9,126 | 9,166 | 9,139 | 9,166 | 9,155 |
| Aligned Bases Ref | 100% | 100% | 100% | 100% | 100% | 100% | 99.98% | 99.98% | 100% | 100% |
| Aligned Bases Query | 100% | 100% | 100% | 100% | 100% | 100% | 99.98% | 99.98% | 100% | 100% |
| Avg Identity | 96.04 | 97.17 | 97.39 | 99.19 | 95.81 | 96.70 | 93.82 | 96.41 | 94.81 | 95.69 |
| Total SNPs | 278 | 192 | 180 | 75 | 318 | 259 | 419 | 226 | 394 | 330 |
| Total Indels | 83 | 58 | 11 | 1 | 81 | 55 | 72 | 9 | 56 | 33 |

### Assembly of simulated viral and bacterial datasets

We tested MetaCortex’s performance on two simulated communities, one viral and one bacterial, both with a high amount of strain level variation and highly variable compositions. For both communities we used the software CAMISIM [25] to simulate the community composition, strain level variants, and 15 Gb of Illumina 2×150bp paired end reads, with a HiSeq 2500 error profile. The simulated viral reads had variable but very high coverage, with coverage ranging from 44,370x to 143,673x for individual taxa across 10 genomes. Previous studies have suggested that the ability of assembly tools to deal with ultra-high coverage genomes is an important but often under-appreciated aspect of virome analysis, particularly when using library preparation methods that increase overall sequencing depth in order to improve recovery of low abundance genomes [28]. The simulated bacterial reads had variable coverage, with coverage ranging from 0.38x to 2,557x for individual taxa.

Using the reference genomes (both real and simulated) we were able to compare the performance of MetaCortex, MEGAHIT, Ray Meta, metaSPAdes and MetaVelvet, on these datasets. Since we assembled the simulated reads directly (i.e., without adapter trimming or other pre-processing), we used the paired-end assembly mode for Ray Meta and MEGAHIT.

The viral community consisted of six species: Human mastadenovirus F, Human herpesvirus 5, Human respiratory syncytial virus, Influenza B virus, Reovirus 3 and Zika virus; and four simulated strains of Influenza B virus. Each simulated strain had between 99.93% and 99.95% of bases aligned to the genome it was simulated from, with an average alignment identity of between 97.09% and 99.63%. The composition of the community is shown in Figure 5.A, and we simulated 150bp long paired-end reads for a total of 14.7Gbp. For this dataset, MetaCortex recovered the highest overall genome fraction (98.78%), with individual genome fractions ranging from 95.27% to 99.91% and had no misassemblies. MEGAHIT recovered a similar genome fraction but had significantly more misassemblies and a higher mismatch rate. Both Ray Meta and metaSPAdes recovered smaller genome fractions (77-93%), although with lower error rates (Table 5). Individual genome coverage for each assembler is displayed in Figure 5.B. MetaCortex achieved the highest NGA50 for the simulated strains (which were also the least abundant) but had lower NGA50 values for some of the other species (Figure 5.C). The assemblies by MetaVelvet either failed to complete, or recovered an insignificant genome fraction, and so are not reported.

**Table 5:**
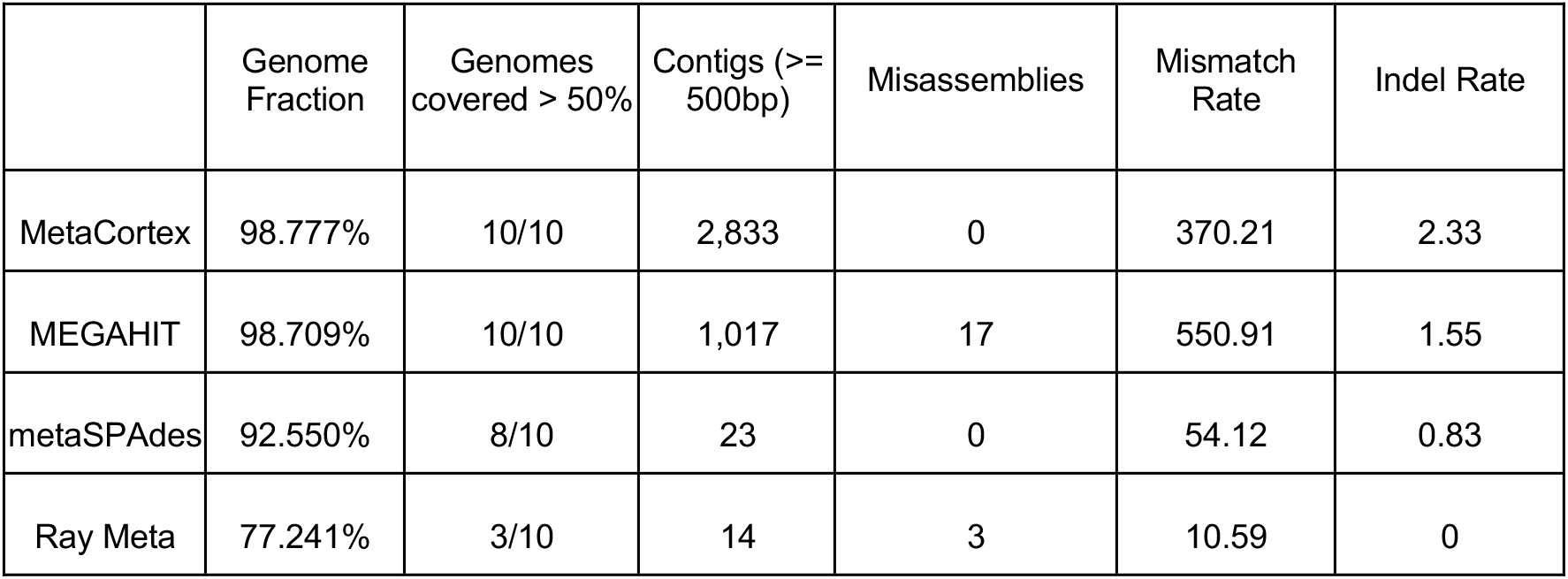
Summary statistics for assemblies of simulated viral community.

**Table 6:**
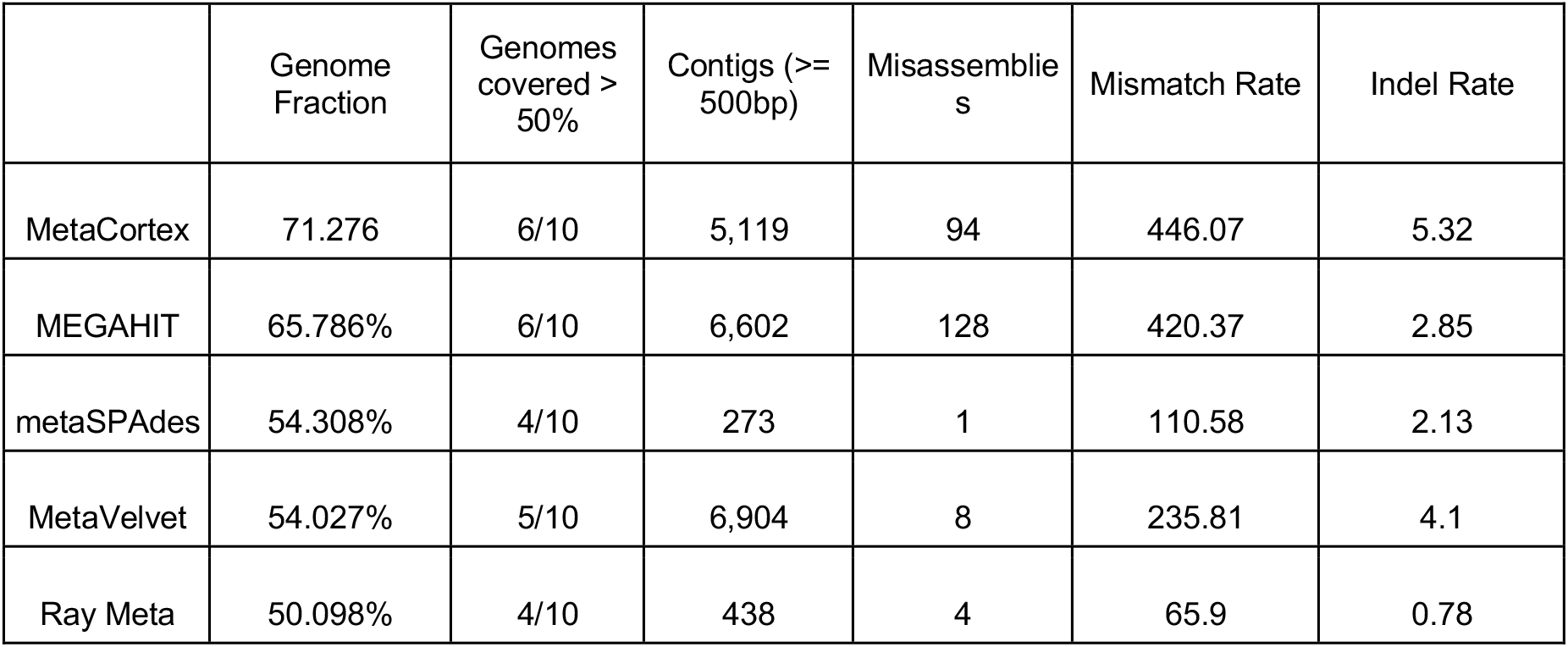
Summary statistics for assemblies of simulated bacterial community.

|  | Genome Fraction | Genomes covered > 50% | Contigs (>= 500bp) | Misassemblies | Mismatch Rate | Indel Rate |
| --- | --- | --- | --- | --- | --- | --- |
| MetaCortex | 71.276 | 6/10 | 5,119 | 94 | 446.07 | 5.32 |
| MEGAHIT | 65.786% | 6/10 | 6,602 | 128 | 420.37 | 2.85 |
| metaSPAdes | 54.308% | 4/10 | 273 | 1 | 110.58 | 2.13 |
| MetaVelvet | 54.027% | 5/10 | 6,904 | 8 | 235.81 | 4.1 |
| Ray Meta | 50.098% | 4/10 | 438 | 4 | 65.9 | 0.78 |

The bacterial community consisted of four species randomly chosen from the well-known MBARC-26 mock community [26]: *Terriglobus roseus*, *Salmonella bongori*, *Fervidobacterium pennivorans*, and *Sediminispirochaeta smaragdinae*; plus two strains simulated from *S. bongori*, and four strains simulated from *F. pennivorans*. Each simulated strain had between 99.98% and 100.00% of bases aligning to the genome it was simulated from, with an average alignment identity of between 97.76% and 99.44%. Using CAMISIM, we simulated 14.9Gbp of 150bp paired-end reads, with the abundances as shown in Figure 5.A.

MetaCortex again recovers the highest genome fraction (71%), with six out of ten genomes recovered at at least 50%. This time, the assembly by MEGAHIT contained the most misassemblies, with MetaCortex containing significantly fewer, but still a high misassembly rate. The assemblies by metaSPAdes, MetaVelvet and Ray Meta contained far fewer misassemblies, but also assembled a much smaller proportion of the genomes present in the sample (27-54%). The assembly by MetaVelvet has the lowest error rates, with no misassemblies, but fails to assemble any sequence for six of the genomes in the community, and for two of the genomes assembles less than 5%. The assembly by metaSPAdes also has low error rates but recovers significantly less of the genomes than MetaCortex, failing to assemble any of the least two abundant strains (F. pennivorans.1 and F. pennivorans.2) and less than 7% of both F. pennivorans.3 and S. bongori.1.

## ALGORITHM

The main innovation that MetaCortex introduces is two new graph traversal algorithms for metagenomic datasets. The core algorithm, MetaCortex Consensus (MC) is able to produce FASTA, GFA, and FASTG outputs. This is the option that should be selected to obtain sequence graphs. The Subtractive Walk algorithm (SW) produces only FASTA output, but performs much faster than MC, and attempts to write a single sequence for each variant that is detected. The user can also choose to only write unitigs (the maximal paths where each inner node has degree 2) as the FASTA output, by specifying the (existing) PerfectPath algorithm.

First, reads are decomposed into *k*-mers, and a de Bruijn graph is constructed; this graph (which will likely consist of several disconnected subgraphs) can be initially pruned to remove nodes which form short ‘tips’, or which fail to meet a minimum level of coverage. Tips up to a length of 100 are pruned by default, and the default minimum coverage is 2 (these values can be modified with command line options). Following construction of the de Bruijn graph, one of the following metagenomic traversal algorithms is executed. Formal descriptions of the algorithms can be found in the Supplementary Materials.

### MetaCortex Consensus algorithm

In a metagenomic read set, disparate species whose genomes do not share any *k*-mers are represented by distinct connected components in the de bruijn graph. This algorithm seeks to find consensus paths through each connected component of the graph, representing the disparate species in the sample, and then represent the inter-species diversity by looking for local topological variation (e.g. bubbles) along each consensus path, and outputting this in sequence graph format (GFA, FASTQ). Thus, the final output is a sequence graph that represents several related taxa.

For each node in the graph, the connected component containing this node is explored to find the highest coverage node and to determine the size of the component. If it is sufficiently large, it is traversed starting at the highest coverage node. At each branching point in the graph, the branch with the highest coverage is favoured if it meets a minimum coverage threshold (Fig. 2a). However, should the paths from two other branches later join together (so that they form a bubble) and have higher coverage collectively, then the highest coverage of these two branches is chosen (Fig. 2b). Traversal continues until a tip is reached, the highest coverage branch at a branching point has already been visited (e.g. in repeat regions), or there are no branches of sufficiently high coverage.

Once a path has been identified, the sequence it represents is written out to a FASTA file. Coverage statistics for the path are included in the header line. If the user has selected to have GFA2/FASTG output, the path is traversed to identify polymorphisms. At each node along the path, any branches that meet a minimum coverage threshold (and are not part of the original path) are explored, depth-first, with the highest coverage node taken at any subsequence branches. If at any point we return to a node on the original path at a position after the original branch, then this new path is written as an alternative path in the GFA file. The path traversal then continues from where the alternative path joins the original path.

Next, each node from the connected component is removed from the graph, and a new connected component is explored. Thus, we obtain one contig and one sequence graph for each sufficiently large connected component. However, if the ‘-M’ flag is specified, only those nodes in the path are removed, the remaining nodes in the current component will be reconsidered, and we obtain multiple contigs for each connected component.

### Subtractive Walk algorithm

One of the key difficulties of metagenomic assembly is the presence of multiple distinct strains (or even distinct species) whose genomes share *k*-mers. This means that in the corresponding de Bruijn graph, some paths may represent portions of the genomes belonging to multiple strains, so it may be desirable to include these in multiple output sequences. The SW algorithm addresses this by not simply removing nodes that have already been traversed, but instead reducing their coverage, so that they may be traversed multiple times.

The algorithm proceeds as follows. First, each node is examined, and for any that meet a minimum coverage requirement, the connected component they are contained in is explored to find the node with locally maximal coverage. (Note that unlike in MC, the whole component may not be explored, in order to speed up the process.) From this node, the highest coverage path is obtained and written to the FASTA file, as in MC, except that at branches, the highest coverage branch is always taken (i.e. bubbles are disregarded).

Next, MetaCortex estimates the number of variants covering each node in the path. First, the lowest coverage node along the path is found and this is assumed to have one variant covering it. There may be more than one node with the same minimal coverage, in which case, the node closest to the end of the path is chosen. Then, starting from the minimal coverage node, the path is walked in each direction, and at each step, the quantity *δ* is calculated, where

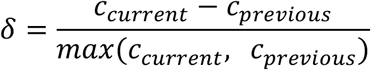

and *c_current_* is the coverage of the node at the current step, and *c_previous_* is the coverage of the node at the previous step. This results in a value between −1 and 1. If this value is less than −*Δ_SW_* (a value determined by the user with option -W, and set to 0.8 by default) then the number of variants covering this node is assigned the value of the number of variants covering the previous node plus 1; if it is greater than *Δ_SW_*, the number of variants covering this node is assigned whichever is larger of the value of the number of variants at the previous node minus 1, and 1.

After assigning a value to each node estimating the number of variants covering that node in the path, the coverage of nodes is adjusted in the following way. Nodes with 1 variant covering them are reduced to 0, and are essentially removed from the graph. Nodes with more than 1 variant covering them are reduced by an amount which is linearly interpolated from the nearest 1-variant nodes before and after this node in the path (Fig. 2c). This process is repeated until all nodes have been examined.

This algorithm is based on the assumption that, for any two adjacent nodes in the graph that represent *k*-mers that only appear consecutively in the metagenome (i.e. the first node has outdegree one, and the second node has indegree one), the change in coverage between them will be small compared to their coverage values. On the other hand, for any two adjacent nodes in the graph that represent *k*-mers that appear consecutively in the metagenome, but at least one of which also appears elsewhere (i.e. at least one of the outdegree of the first node and indegree of the second node is greater than one), the change in coverage between them may be significant compared to the coverage values. The relative change in coverage across the path is what the value *δ* measures. This assumption, however, is only true for samples that have been sequenced with shotgun metagenome sequencing, and for regions of low sequence complexity (for example, repeat regions) this may not be the case.

When choosing a value for the parameter *Δ_SW_*, the user should choose smaller values if the dataset is expected to have high levels of strain diversity and the user wishes to capture this in the assembly. For datasets with less strain diversity, or if the user wishes to capture only dominant strains, higher values (closer to 1.0) should be chosen.

## IMPLEMENTATION

MetaCortex uses Cortex’s hash table structure to store *k*-mer information and to encapsulate the de Bruijn graph structure. For reasons of memory efficiency, the maximum *k*-mer size must be specified when building MetaCortex. The default maximum value is 31, with 63, 95,127,160 or 192 also possible. The size of the hash table is user-defined and should be sufficient to contain the totality of the dataset being assembled (if when loading the reads into the hash table, it becomes full, the user is warned, but execution continues and no new *k*-mers are added to the hash table). Further details can be found in MetaCortex’s documentation and the cortex_var manual (http://cortexassembler.sourceforge.net/cortex_var_user_manual.pdf, accessed November 2021).

After the construction of the hash table from the read files, a binary representation of the de Bruijn graph can be written to disc and this can then be used as the input to later assemblies. This can be used to speed up assembly time for subsequent assemblies of the same dataset, or to parallelise the reading of FASTA or FASTQ files. In the latter case, the individual CTX files can be merged to construct a de Bruijn graph from the whole read set. Figure 1 depicts a typical workflow.

**Figure 1:**
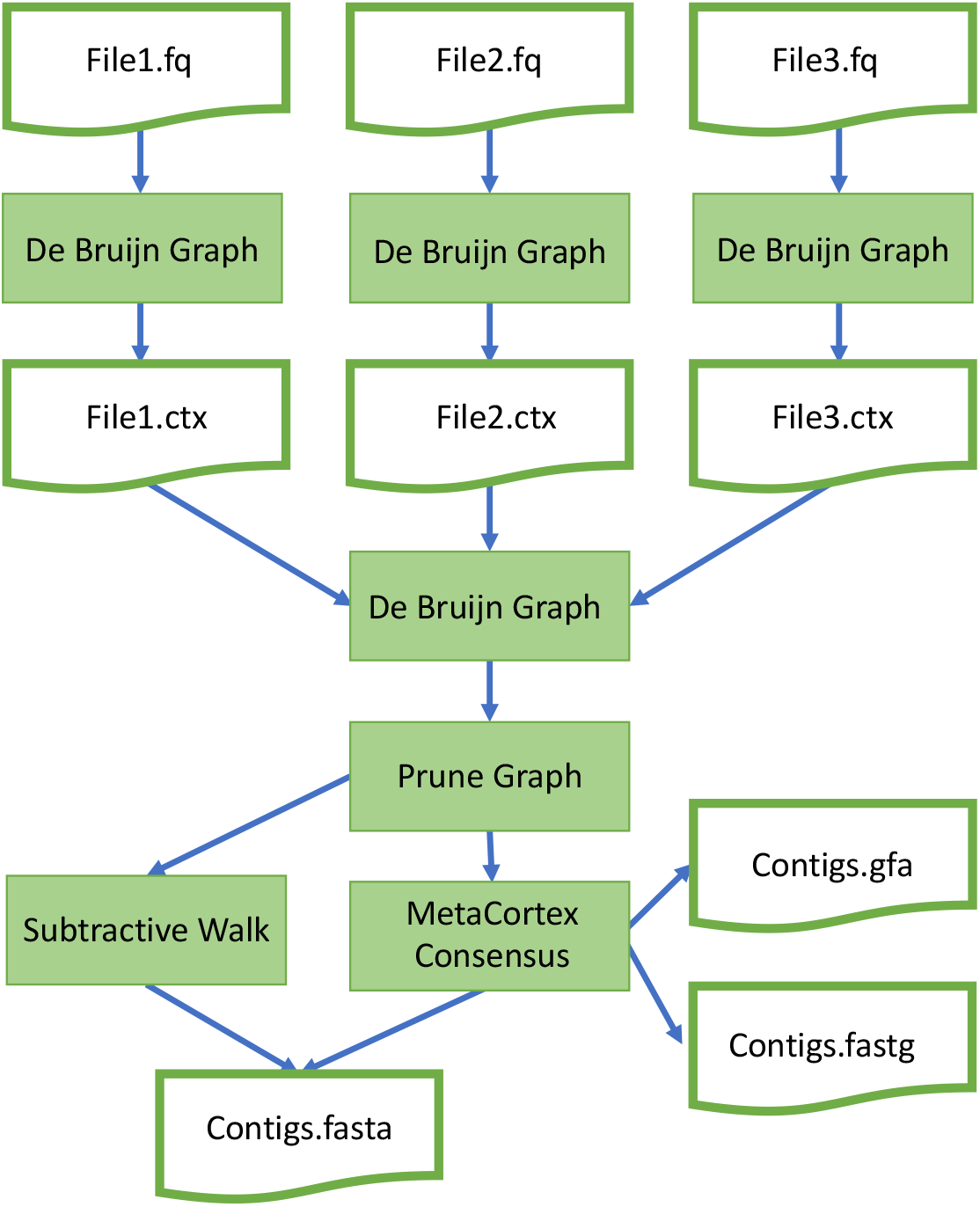
Flow chart depicting typical MetaCortex workflow.

**Figure 2:**
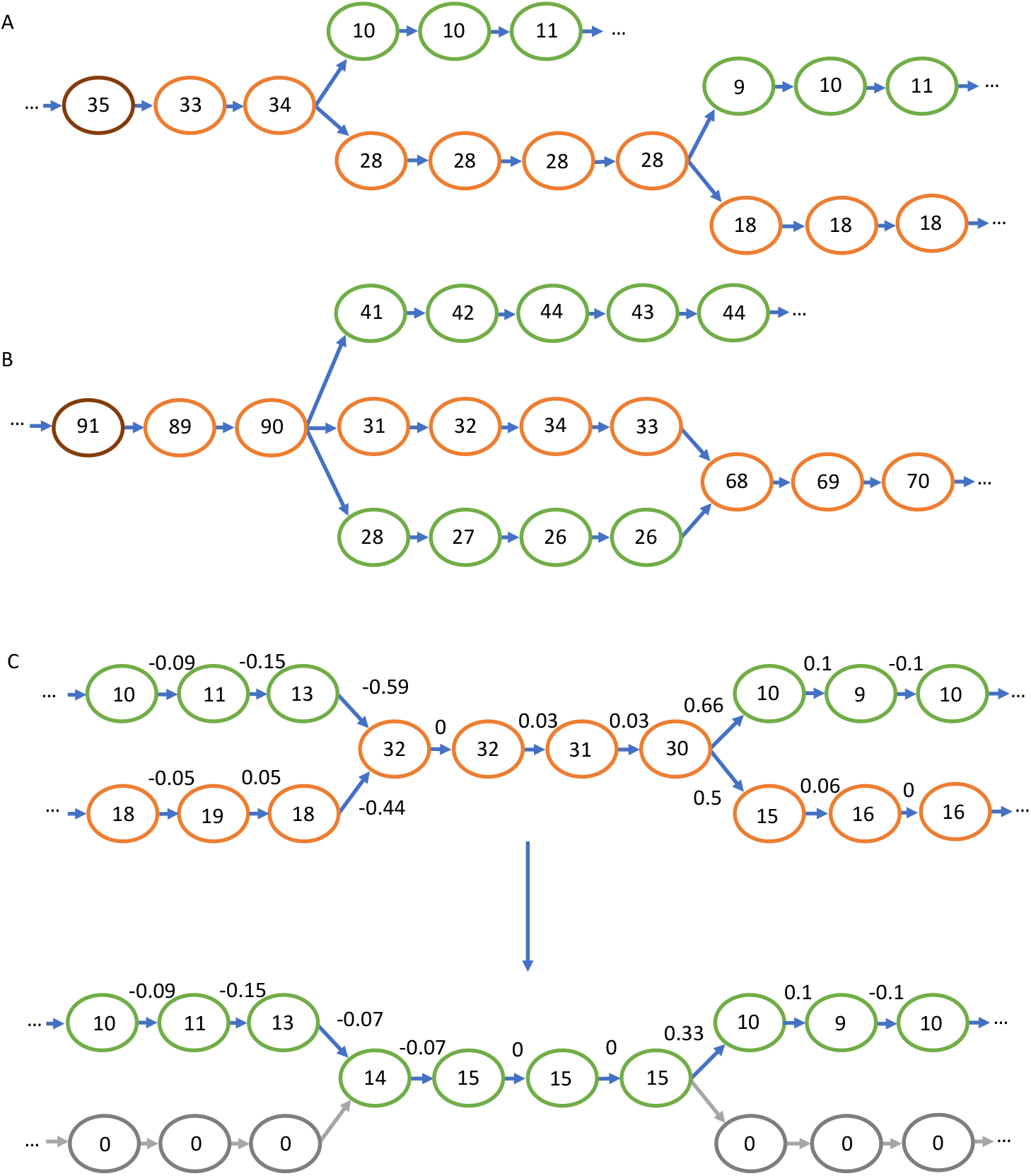
Depictions on de Bruijn graphs, with the coverage for each node represented. **A**. The path chosen by MetaCortex Consensus, with the highest coverage node highlighted in brown, and the chosen path in orange. **B**. The path chosen when two of the branches form a bubble. Because the two bubble branches, added together, represent a combined higher coverage than the top branch, a route through the bubble is selected for the path. **C**. The progression of the Subtractive Walk algorithm. The numbers above the edges are the normalised coverage difference *δ*. The first graph is before coverage subtraction, with the path chosen highlighted in orange. The second graph is after coverage subtraction. Nodes belonging to exactly one path are removed (shown in grey), whilst nodes that are shared between paths remain with reduced coverage.

**Figure 3:**
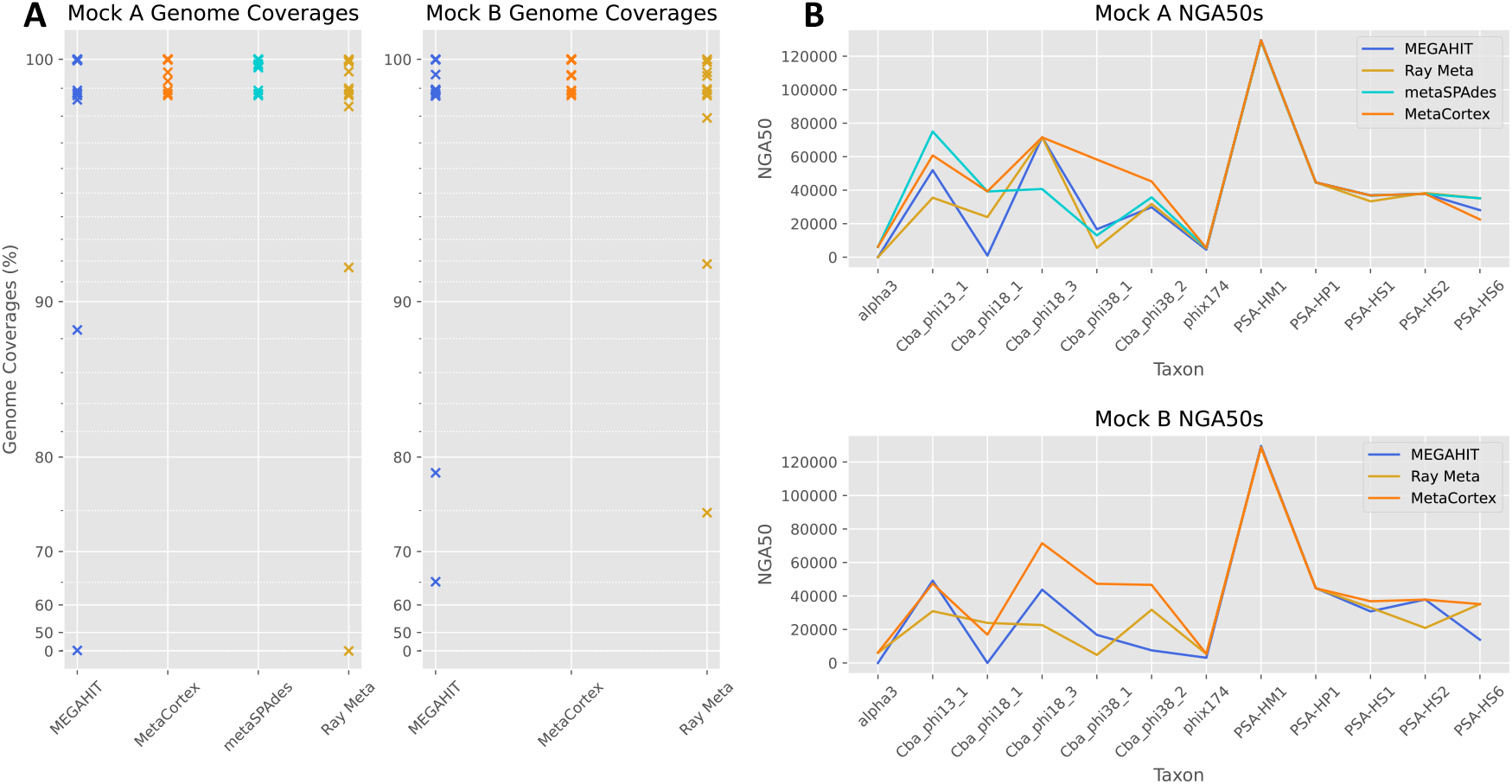
**A**. Genome coverages for Mock A and Mock B for assemblies presented in Table 1 and Table 2. **B**. NGA50 by viral species for assemblies presented in Table 1 and Table 2.

**Figure 4.**
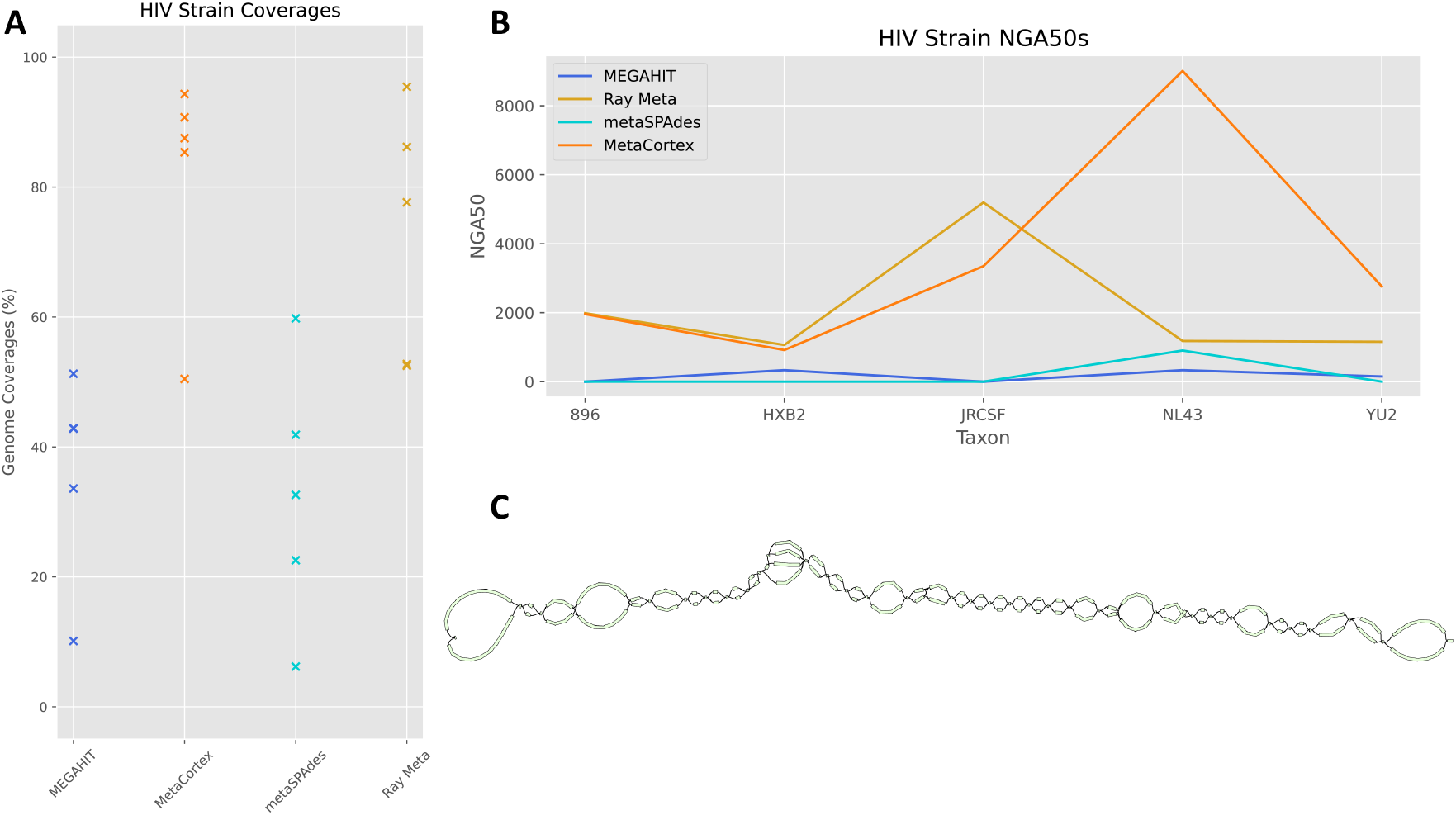
Assemblies of HIV-5-Strain dataset. **A**. Coverage for each viral species for each assembler. **B**. NGA50s for each strain per assembler. **C**. Visualisation of GFA sequence graph output from MetaCortex. The sequence graph shows a single contig, with local variation shown by branching sequences.

**Figure 5:**
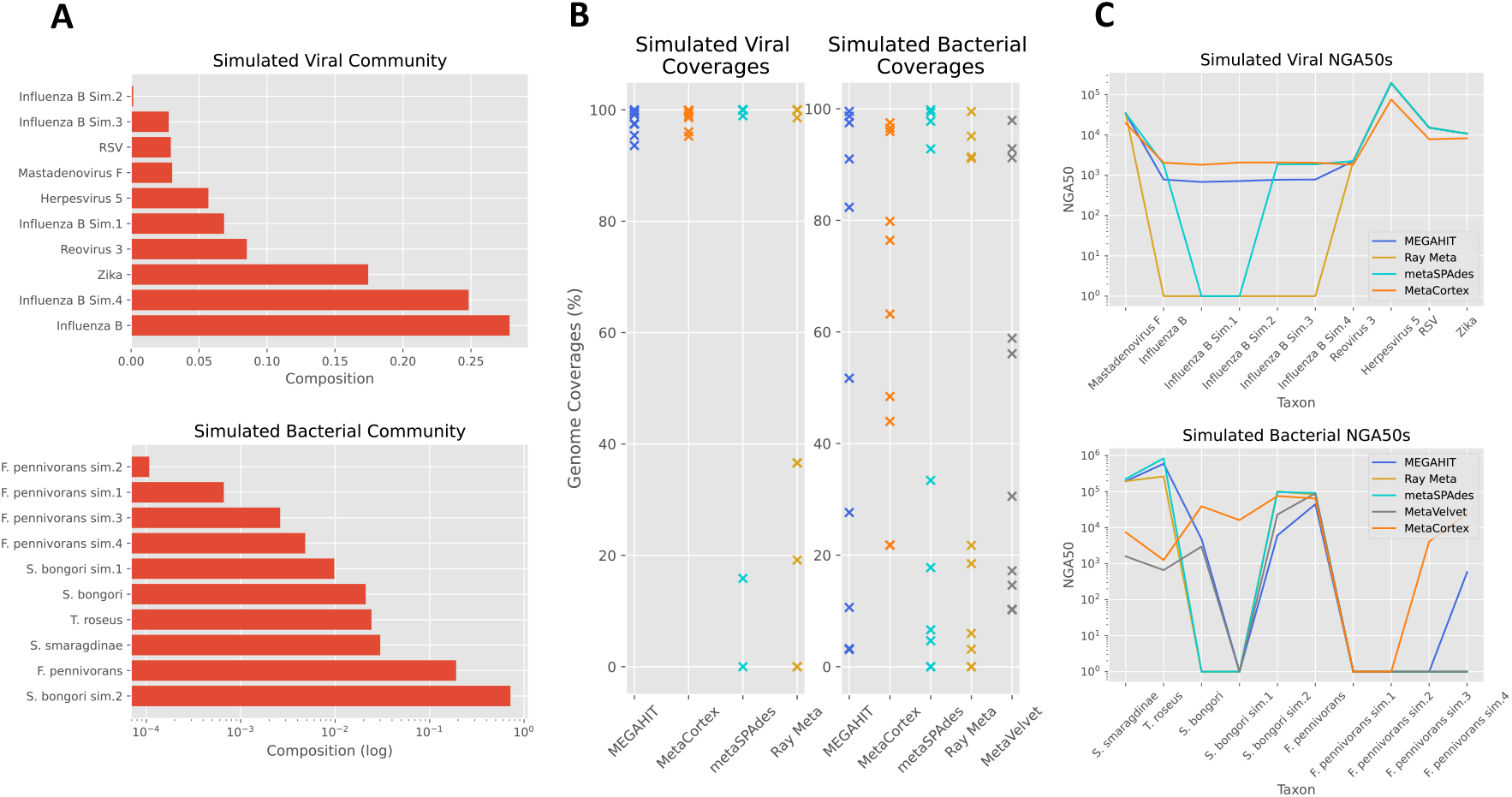
**A**. Compositions of simulated communities. **B**. Coverage per species for each assembler. **C**. NGA50s of each species by assembler (log scale).

## DISCUSSION

Compared to the challenge of assembling a single isolate species, metagenomic assembly presents significant additional hurdles related to the presence of closely related species and their differing abundance in a sample. These challenges can be particularly obvious in viral communities, where a rapid evolutionary rate can make it especially difficult to distinguish between and within quasispecies. In recent years, several new assembly tools have attempted to tackle the challenges of metagenomic assembly using various heuristic approaches. To our knowledge, all have adopted approaches which reduce assembly graphs to sets of contigs, losing the variation captured by the underlying graph structures. We address this in MetaCortex, a new assembly tool that preserves strain interconnectedness by outputting sequence graph files as an alternative to contigs. In addition, a new Subtractive Walk algorithm enables MetaCortex to estimate the number of variants in each subgraph and to output representative contigs. This algorithm was able to recover a high proportion of five strains of HIV from a lab mixed community within a metagenomic dataset, and five strains (one real, four simulated) of influenza virus from a simulated viral community, without misassembly. The latter result is particularly encouraging, as several of these strains were at very low abundance in the sample.

Overall, we found that MetaCortex consistently recovers a very high genome fraction when compared to other popular metagenome assemblers. In particular, our simulated datasets show that MetaCortex is especially effective at recovering the genomes of extremely low abundant species. For the assembly of viral communities, MetaCortex had low error rates comparable with the lowest of the other assemblers tested.

## DATA AVAILABILITY

MetaCortex is open source and available to download from https://github.com/SR-Martin/metacortex. Full documentation, including instructions for installation, is available at https://metacortex.readthedocs.io/en/latest/. All assemblies, simulated reads, and simulated genomes used in this paper have been deposited online on Zenodo and can be found at DOI 10.5281/zenodo.6616437.

## SUPPLEMENTARY DATA

Supplementary Data are available online.

## ACKNOWLEDGEMENTS

We are grateful to Kirsten McLay for helping to enable some of the laboratory work that led to this publication. This research was supported in part by the NBI Computing Infrastructure for Science Group, which provides technical support and maintenance to Earlham Institute’s high-performance computing cluster and storage systems.

