## Supplementary Materials for "Capturing variation in metagenomic assembly graphs with MetaCortex"

### 1 Algorithm Description

In this section we give pseudocode for the two main graph traversal algorithms employed by MetaCortex. We use the convention that the de Bruijn graph constructed by MetaCortex is a digraph $G$, with vertex set $V$ given by the distinct $k$-mers, and edge set $E \subseteq V \times V$, where an edge $e = (v_1, v_2)$ is in $E$ if and only if the subsequence of length $k-1$ at the end of the $k$-mer associated with $v_1$ is equal to the subsequence of length $k-1$ at the beginning of the $k$-mer associated to $v_2$. An edge $(v_1, v_2)$ therefore has a letter associated to it, given by the final letter of the $k$-mer at $v_2$. For a $k$-mer represented by a vertex $v$, the vertex associated to the reverse complement of this $k$-mer will be denoted $v^c$. Each vertex $v$ also has a non-negative integer associated with it, which describes the number of times the $k$-mer represented by $v$ appears in the read set. We have a function *Coverage*, which given a vertex $v \in V$, returns the coverage of $v$. When a vertex has 0 coverage, we consider it, and all edges incident to it, as removed from the graph.

Contiguous sequences of DNA can be constructed from paths in the graph. Paths in the graph are represented by finite sequences of vertices $(v_0, v_1, \ldots, v_n)$ such that for $i = 0, \ldots, n-1$ we have $(v_i, v_{i+1}) \in E$. The sequence given by this path consists of the $k$-mer at $v_0$, followed by the letters associated to each subsequent edge in the path. For $v \in V$, the connected component of $G$ containing $v$ is the subgraph of $G$ with vertex set $W$ consisting of all vertices $w \in V$ such that there exists a path between $v$ and $w$, and edge set given by $E \cap (W \times W)$. We have two functions on paths. First, *Concatenate*, which given two paths $P_1 = (v_1, \ldots, v_n)$ and $P_2 = (w_1, \ldots, w_m)$ with $v_n = w_1$, returns the path $(v_1, \ldots, v_n, w_2, \ldots, w_m)$. Second, *ReverseComplement*, which, given a path $P = (v_1, \ldots, v_n)$, returns the path $(v_n^c, \ldots, v_1^c)$.

**Algorithm 1:** Algorithm for finding the highest coverage path from a starting node.

function HighestCoveragePath $(v_0, G, C_{\min})$;
**Input :** Starting node $v_0$, de Bruijn graph $G = (V, E)$, and minimum node coverage $C_{\min}$
**Output:** Path $P$
Path $P \leftarrow (v_0)$;
Node $v \leftarrow v_0$;
**while** $w \in V$ *such that* $(v, w) \in E$*, $w$ is unvisited, and Coverage*$(w)$ *is maximal among all nodes* $w'$ *such that* $(v, w') \in E$ **do**
  **if** *Coverage*$(w) \geq C_{min}$ **then**
    append $w$ to $P$;
    mark $w$ and $w^c$ as visited;
    $v \leftarrow w$;
  **else**
    return $P$;
  **end**
**end**
return $P$;

**Algorithm 2:** Algorithm for finding the highest coverage path from a starting node, preferring bubbles over single paths when the combined coverage of the bubble is larger than that of any single path.

```
function HighestCoverageBubblePath ($v_0, G, C_{\min}$);
Input  : Starting node $v_0$, de Bruijn graph $G = (V, E)$, and minimum node coverage $C_{\min}$
Output: Path $P$
Path $P \leftarrow (v_0)$;
Node $v \leftarrow v_0$;
while $w \in V$ such that $(v, w) \in E$, $w$ is unvisited, and Coverage$(w)$ is maximal among all
  nodes $w'$ such that $(v, w') \in E$ do
    Int $c_{\max} \leftarrow C_{\min}$;
    Node $v_{\text{next}} \leftarrow \varnothing$;
    if Coverage$(w) > c_{max}$ then
        $c_{\max} \leftarrow$ Coverage$(w)$;
        $v_{\text{next}} \leftarrow w$;
    end
    foreach pair $(w_1, u_1) \in V \times V$ with $(v, w_1), (v, u_1) \in E$ and $w_1, u_1$ unvisited do
        Let $P_w = (w_1, \ldots, w_n)$ be the path from $w_1$ with $\deg^+(w_i) = 1$ for $i = 1, \ldots, n-1$ and
          $\deg^+(w_n) > 1$;
        Let $P_u = (u_1, \ldots, u_m)$ be the path from $u_1$ with $\deg^+(u_i) = 1$ for $i = 1, \ldots, m-1$ and
          $\deg^+(u_m) > 1$;
        if $w_n = u_m$ then
            $c \leftarrow$ Coverage$(w_1)$ + Coverage$(u_1)$;
            if $c > c_{max}$ then
                $c_{\max} \leftarrow c$;
                $v_{\text{next}} \leftarrow$ the node in $\{w_1, u_1\}$ with highest coverage;
            end
        end
    end
    if $v_{next} \neq \varnothing$ then
        append $v_{\text{next}}$ to $P$;
        mark $v_{\text{next}}$ and $v^c_{\text{next}}$ as visited;
        $v \leftarrow v_{\text{next}}$;
    else
        return $P$;
    end
end
return $P$;
```

**Algorithm 3:** MetaCortex Consensus algorithm.

function MetaCortexConsensus ($G, C_{\min}, m$);<br>
**Input :** De Bruijn graph $G = (V, E)$, minimum node coverage $C_{\min}$, and boolean value $m$ indicating whether only one path per connected subgraph should be returned.<br>
**Output:** Array of paths $\mathcal{P}$<br>
Array $\mathcal{P} \leftarrow \varnothing$;<br>
**foreach** *node* $v \in V$ **do**<br>
    **if** $v$ *is unvisited* **then**<br>
        Let $S$ be the connected component of $G$ containing $v$;<br>
        Find the node $v_{\text{max}}$ in $S$ with maximal coverage;<br>
        $P_{\text{forward}} \leftarrow$ HighestCoverageBubblePath($v_{\text{max}}, G, C_{\min}$);<br>
        $P_{\text{reverse}} \leftarrow$ HighestCoverageBubblePath($v^c_{\text{max}}, G, C_{\min}$);<br>
        $P \leftarrow$ Concatenate(ReverseComplement($P_{\text{reverse}}$), $P_{\text{forward}}$);<br>
        **if** $P \neq \varnothing$ **then**<br>
            Append $P$ to $\mathcal{P}$;<br>
            **if** $m$ = *True* **then**<br>
                Remove $S$ from $G$;<br>
            **else**<br>
                Remove $P$ from $G$;<br>
            **end**<br>
        **end**<br>
    **end**<br>
**end**<br>
return $\mathcal{P}$;

**Algorithm 4:** Subtractive Walk algorithm.

function SubtractiveWalk $(G, C_{\min}, \Delta_{\min})$;
**Input :** De Bruijn graph $G = (V, E)$, minimum node coverage $C_{\min}$, and SW-delta $\Delta_{\min}$
**Output:** Array of paths $\mathcal{P}$
Array $\mathcal{P} \leftarrow \varnothing$;
**foreach** *node* $v \in V$ **do**
  Explore the graph $G$ around $v$ to find the node $v_0$ with locally highest coverage;
  **if** *Coverage*$(v_0) > C_{min}$ **then**
    $P_{\text{forward}} \leftarrow \text{HighestCoveragePath}(v_{\max}, G, C_{\min})$;
    $P_{\text{reverse}} \leftarrow \text{HighestCoveragePath}(v^c_{\max}, G, C_{\min})$;
    $P \leftarrow \text{Concatenate}(\text{ReverseComplement}(P_{\text{reverse}}), P_{\text{forward}})$;
    Append $P$ to $\mathcal{P}$;
    Let $C[]$ be an integer array indexed by the nodes of $P$;
    Let $w_0$ be the node in $P$ with minimal coverage and closest to the end of $P$;
    Int $C_{\text{current}} \leftarrow 1$;
    $C[w_0] \leftarrow C_{\text{current}}$;
    **foreach** *node* $w \in P$ *starting from the node after* $w_0$ *until the last node in* $P$ **do**
      Let $w_{\text{prev}}$ be the node in $P$ before $w$;
      Float $\Delta \leftarrow \frac{\text{Coverage}(w) - \text{Coverage}(w_{\text{prev}})}{\max(\text{Coverage}(w), \text{Coverage}(w_{\text{prev}}))}$;
      **if** $\Delta > \Delta_{min}$ **then**
        $C_{\text{current}} \leftarrow C_{\text{current}} + 1$;
      **else if** $\Delta < -\Delta_{min}$ **then**
        $C_{\text{current}} \leftarrow \max(1, C_{\text{current}} - 1)$;
      $C[w] \leftarrow C_{\text{current}}$;
    **end**
    $C_{\text{current}} \leftarrow 1$;
    **foreach** *node* $w \in P$ *starting from the node before* $w_0$ *until the first node in* $P$ **do**
      Let $w_{\text{next}}$ be the node in $P$ after $w$;
      Float $\Delta \leftarrow \frac{\text{Coverage}(w) - \text{Coverage}(w_{\text{next}})}{\max(\text{Coverage}(w), \text{Coverage}(w_{\text{next}}))}$;
      **if** $\Delta > \Delta_{min}$ **then**
        $C_{\text{current}} \leftarrow C_{\text{current}} + 1$;
      **else if** $\Delta < -\Delta_{min}$ **then**
        $C_{\text{current}} \leftarrow \max(1, C_{\text{current}} - 1)$;
      $C[w] \leftarrow C_{\text{current}}$;
    **end**
    **foreach** *node* $w \in P$ **do**
      **if** $C[w] = 1$ **then**
        Reduce Coverage$(w)$ to 0;
      **else**
        Decrease Coverage$(w)$ by an amount linearly interpolated between nearest two nodes $w_1$ and $w_2$ with $w_1 < w < w_2$ and $C[w_1], C[w_2] > 1$;
      **end**
    **end**
  **end**
**end**
return $\mathcal{P}$;

### 2 Commands

#### 2.1 Metagenomic Assemblies

In this section we give the commands used to obtain the results in the main paper. Reads were trimmed using Trim Galore v0.5.0 and the following commands. Since MetaCortex does make use of paired end information, but other assemblers do, we trimmed each real read set twice - once treating all reads as single ends, and once using paired-end information. For assemblers that can assemble both single end and paired end reads, we assembled each dataset twice.

Listing 1: Read trimming commands

```
#!/bin/bash
trim_galore  all_reads.fq
trim_galore  --paired --retain_unpaired all_R1.fq all_R2.fq
```

The following commands were used to perform the assemblies.

Listing 2: MetaCortex command

```
#!/bin/bash
for k in 31 63 95 127; do
        echo all_reads_trimmed.fq > reads.txt
        mkdir MetaCortex_k${k}
        metacortex_k127-k ${k} -n 28 -b 100 -i reads.txt -t fastq -g 500 \
        -l MetaCortex_k${k}/log.txt -f MetaCortex_k${k}/contigs.fa \
        -C 10 -W 0.8 -A SW
done
```

Listing 3: Ray Meta commands

```
#!/bin/bash
for k in 31 63 95 127; do
        Ray meta -k ${k} -o RayMeta_k${k}_single -s all_reads_trimmed.fq
        Ray meta -k ${k} -o RayMeta_k${k}_paired \
        -p all_R1_val_1.fq all_R2_val_2.fq \
        -s unpaired.fq
done
```

Listing 4: MEGAHIT commands

```
#!/bin/bash
megahit -r all_reads_trimmed.fq -t 8 -o MEGAHIT_single
megahit -1 all_R1_val_1.fq -2 all_R2_val_2.fq -r unpaired.fq -t 8 \
-o MEGAHIT_paired
for m in 5 10 20; do
        megahit -r all_reads_trimmed.fq -t 8 -o MEGAHIT_m${m}_single \
        --min-count ${m}
        megahit -1 all_R1_val_1.fq -2 all_R2_val_2.fq -r unpaired.fq -t 8 \
        -o MEGAHIT_m${m}_paired --min-count ${m}
done
```

Listing 5: metaSPAdes command

```
#!/bin/bash
mkdir metaSPAdes
metaspades.py -1 all_R1_val_1.fq -2 all_R2_val_2.fq -o ./metaSPAdes
```

Listing 6: MetaVelvet commands

```
#!/bin/bash
perl shuffleSequences_fastq.pl R1_val_1.fq R2_val_2.fq interleaved.fq
```

```
for k in 31 63 95; do
        velveth MetaVelvet_k${k} ${k} -shortPaired -fastq interleaved.fq
        velvetg MetaVelvet_k${k}  -exp_cov auto
        meta-velvetg MetaVelvet_k${k}
done
```

Assemblies were assessed against reference sequences with MetaQUAST using the following command.

Listing 7: MetaQUAST command

```
#!/bin/bash
mkdir MetaQUAST
metaquast.py ${assemblies} -o MetaQUAST --ambiguity-usage one \
-R ${references}
```

#### 2.2 GFA Analysis

The strain specific contigs described in 'GFA output captures strain level variation and facilitates visualisation' were produced using the following commands. The script 'strain_assembly.py' is included in the MetaCortex distribution.

Listing 8: Assembly of strain-specific contigs

```
#!/bin/bash
metacortex_k127-k ${k} -n 100 -b 20 -i reads.txt -t fastq\
                 -f MetaCortex_HIV.fa -l log.txt -C 1000\
                 -g 500 -M -G -A MCC
awk '/^S/{print ">"$2"\n"$4}' MetaCortex_HIV.gfa > MetaCortex_HIV.gfa.fa
blastn -db ./references/5-virus-mix.fasta -query MetaCortex_HIV.gfa.fa\
                -task megablast -out MetaCortex_HIV.gfa.blast \
                -outfmt '6 qseqid sseqid pident length qlen'
python ${METACORTEX_DIR}/scripts/strain_assembly.py -g MetaCortex_HIV.gfa\
                 -b MetaCortex_HIV.gfa.blast
mkdir dnadiff_strains
mkdir dnadiff_original

for STRAIN in HXB2 NL43 JRCSF 896 YU2; do
        dnadiff -p dnadiff_strains/${STRAIN} ./references/${STRAIN}.fasta\
                        ./${STRAIN}_gfa_assembled.fa
        dnadiff -p dnadiff_original/${STRAIN} ./references/${STRAIN}.fasta\
                       ./MetaCortex_HIV.fa
done
```
